# Abnormalities of the liver have big impacts on orthotopic liver transplantation

**DOI:** 10.1101/2024.10.12.617996

**Authors:** Yongfeng Chen, Wenzhong Li, Guoyong Chen, Shaotang Zhou

## Abstract

**Background:** Orthotopic rat liver transplantation (OLT) is widely used in basic research; normal liver anatomy and structures are attributable to its success while its deformities complicate to have a negative on OLT.

**Methods:** For tolerance induction project, we performed OLT from Lewis to Brown Norway (BN) rat as chronic rejection model and encountered two anatomical deformities of recipient rats: of 47 liver transplantations, accessory liver lobe occurred to 4 cases and bifurcations of liver outflow to 5 cases in BN rats.

**Results:** For the accessory liver lobe, we discontinued OLT for one case with a big accessory liver lobe; two rats died from pneumothorax upon separation; and succeed in one case with the small lobe. For two vein outflows of the liver, we succeeded in OLT due to its reconstruction in one case but the recipient died one week later, and succeeded in 1 case after one small orifice was sutured, we failed in 3 cases due to thrombosis following OLT. For 38 rats with normal livers, only 4 rats failed to survive LT. There were significant differences in OLT success (*p*<0.01)

**Conclusion:** Liver abnormal anatomy has negative impact on OLT, giving a clue to that pre-transplant comprehensive screening is significant and beneficial clinically.

## Introduction

Rat OLT receives wide acceptance as a small animal model in the research of ischemia-reperfusion injury, immunology and transplant medicine, especially in tolerance induction ever since its development by Lee and his colleagues [1,2], it is a complicated operation requiring good microsurgical performance, normal liver anatomy and condition etc [3-6]. Here we first reported the anatomical deformities of the rat liver which have negative impact on OLT.

## Methods

We are studying liver regeneration and immunological tolerance through stem cells (detailed protocol out of scope here) for our project and perform rat OLT from Lewis to BN that it is well recognized as a chronic rejection model [5]. Male Lewis rats (weighing 200-280g) and BN serve as donors and recipients respectively and were purchased from Vital River Laboratory Animal Technology Corporation. The animals were housed and cared in the standard environment with 12 hour-cycle of light and darkness at the controlled temperature, they freely access to a standard food and water; each rat was fasted 12 hours prior to OLT. All experiments were approved by the Ethics Committee of Henan Integrated Traditional Chinese and Western Medicine Hospital and conducted in compliance with the standards for animal use and care set by ARRIVE guidelines and the Institutional Animal Care Committee (No: HNTCMDW-20240829).

### Surgical procedure

Some modifications were made for isoflurane inhalation anesthesia applied to the project. In the donor procedure, one microsurgeon made a transverse incision to enter the abdominal cavity of the rat, he retrogradely flushed the donor liver first through the aorta with hepatized normal saline (50 u/ml) and then reflushed through the portal vein (PV) with 4 ^0^C lactated Ringer’s solution. The artery segments from the celiac to the proper liver one was kept; the splenic artery and the left gastric one were ligated respectively. The graft was explanted out and stored in lactated Ringer’s solution. Cuffs were prepared for PV (outer diameter 2.6 mm, inner diameter 2.2 mm) and infra-hepatic vena cava (IVC. outer diameter 3.2 mm, inner diameter 2.8 mm). Cold storage time was less than 3 hours in all cases. In the recipient, a midline incision was made and the surgical field was exposed by 3 self–made retractors with clips; all ligaments of the liver were separated with the electric bipolar. The proper liver artery was ligated; surgeon made a blunt separation behind the liver to create a tunnel. The recipient PV and IVC were clamped sequentially with microvascular clamps, and isoflurane was immediately decreased to 0.2 volume %. Through the tunnel, a mosquito forceps was positioned on the part of diaphragm ring (left side) to occlude the suprahepatic vena cava (SHVC), the native liver was taken out. SHVC was anastomosed with 8-0 polypropylene suture, when completed, the mosquito forceps was displaced with a vascular bulldog on the real SHVC while the diaphragm ring was de-clamped. PV was reconnected with the cuff; blood flow was restored when the clamp on the PV was released, the an-hepatic time was generally less than 20 minutes. IVC reconnection was made as it was for PV. Microsurgeon ligated the gastro-duodenal artery proximally and made an opening on the common hepatic artery into which the stent (self-made) in the donor hepatic artery was inserted and secured [5], bile duct continuity was made when a tube (0.8 mm in outer diameter) in the donor bile duct was inserted into the recipient bile duct. The abdomen was closed with two layers and the animals were placed in a cage under an infrared light. 10% glucose solution and purified water were both supplied for 3 first days, and later regular food and tap water were offered. OLT with recipient alive for 2 days was defined as success.

### Statistical analysis

The success rates of LT in the different groups were evaluated with the X^2^ test which was calculated artificially and P<0.05 was considered significant difference.

## Results

For explant of the recipient liver, we encountered two anatomical abnormalities of BN rats: accessory liver lobe in 4 cases (4/47) and two liver outflows in 5 cases (5/47) (Table 1). The accessory liver lobe was surrounded by ligament connecting the diaphragm and caused Morgagni hernia (Figure 1, 2), one was small accessory lobe which was separated from the main liver using the bipolar cauterizer, and OLT was successful (supplementary video 1). The accessory lobe were bigger in 2 cases, separation was made unsuccessful due to pneumothorax and led to death in one case, it was discarded as the recipient in 1 case. For abnormal outflow of the liver, it occurred in 5 cases (Figure 3). 2 outflows were made septic by muscular diaphragm and converge into one inferior vena cava in the thoracic cavity. OLT was completed due to reconstruction of the outflows in one case and the host died in one week, it succeeded in a case after one small orifice (left) was sutured (Figure 4); OLT failed in 3 cases due to big thrombi shortly after systemic blood restoration. Of remaining 38 rats, 34 underwent OLT successfully and 4 failed due to carelessness. There were significant difference in LT success (*p*<0.01).

**Table 1.** LT and LT with deformities. *X*^***2***^=18.386, *p*<0.01

| Liver | Cases | LT success | LT failure |
| --- | --- | --- | --- |
| accessory lobe | 4 | 1 | 3 |
| outflows | 5 | 1 | 4 |
| normal | 38 | 34 | 4 |

**Figure 1.**
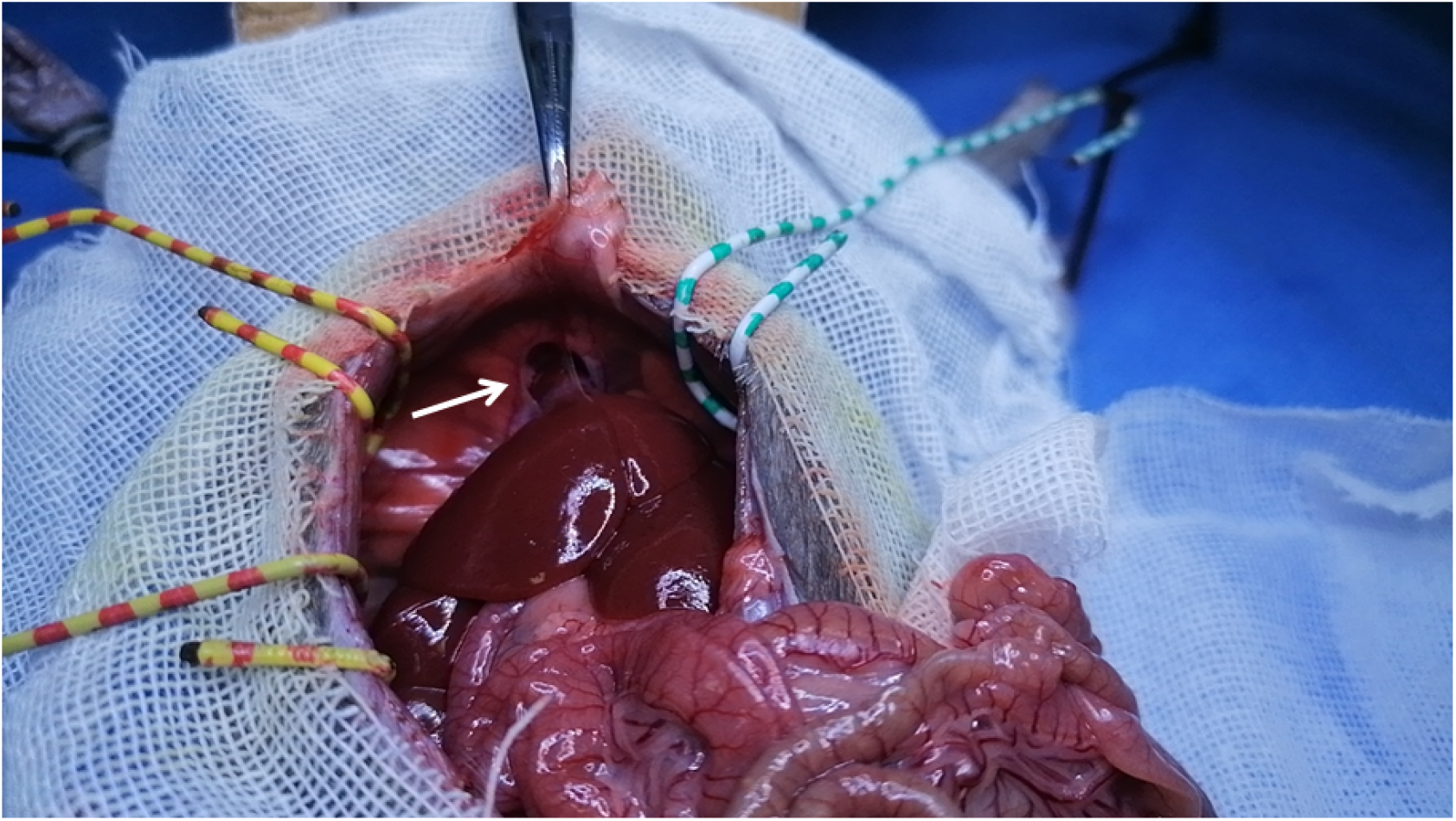
Accessory liver lobe.It is connected to the diaphragm (arrow).

**Figure 2.**
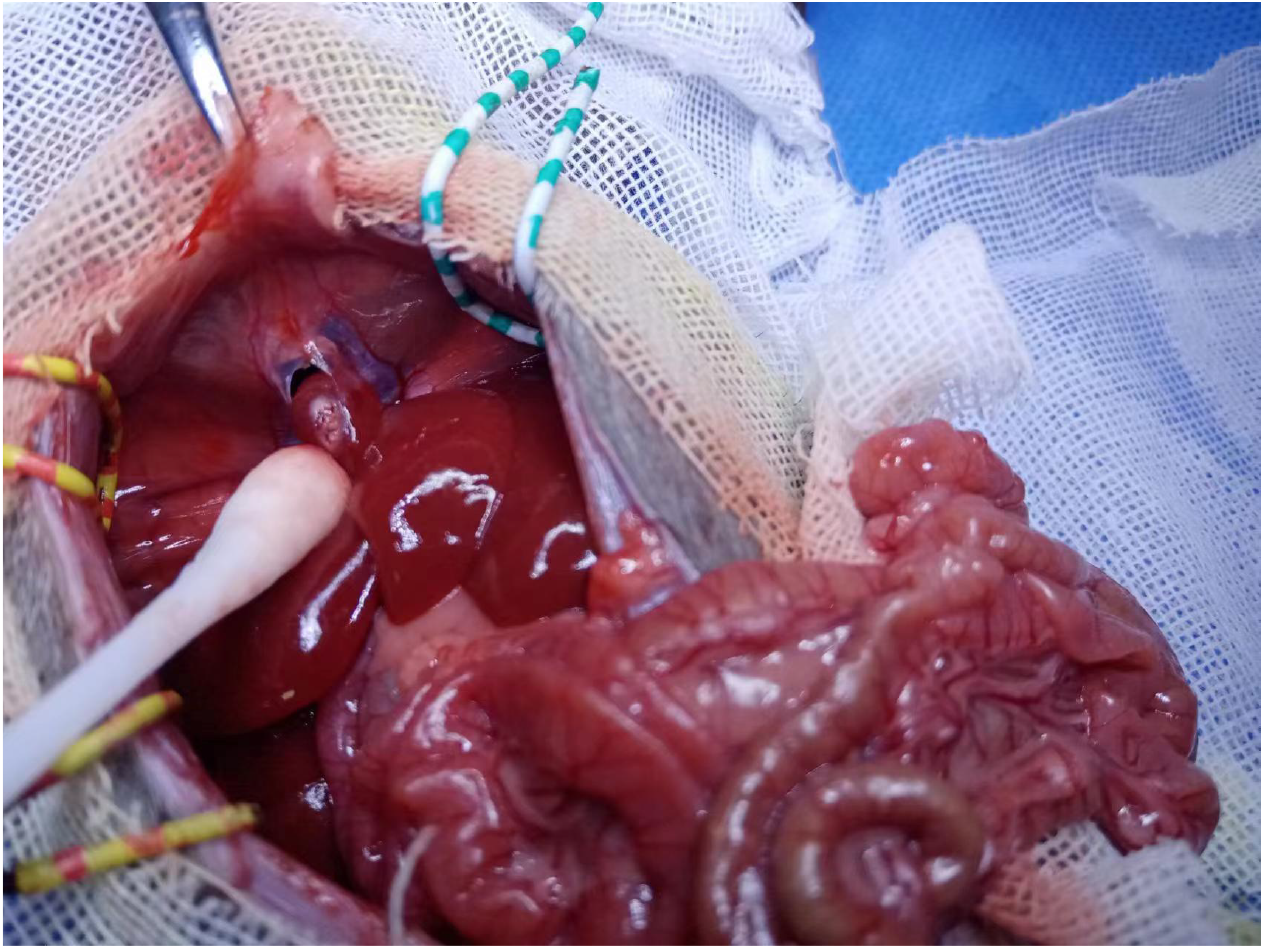
Accessory liver lobe. Pneumothorax occurred and cause death after separation.

**Figure 3.**
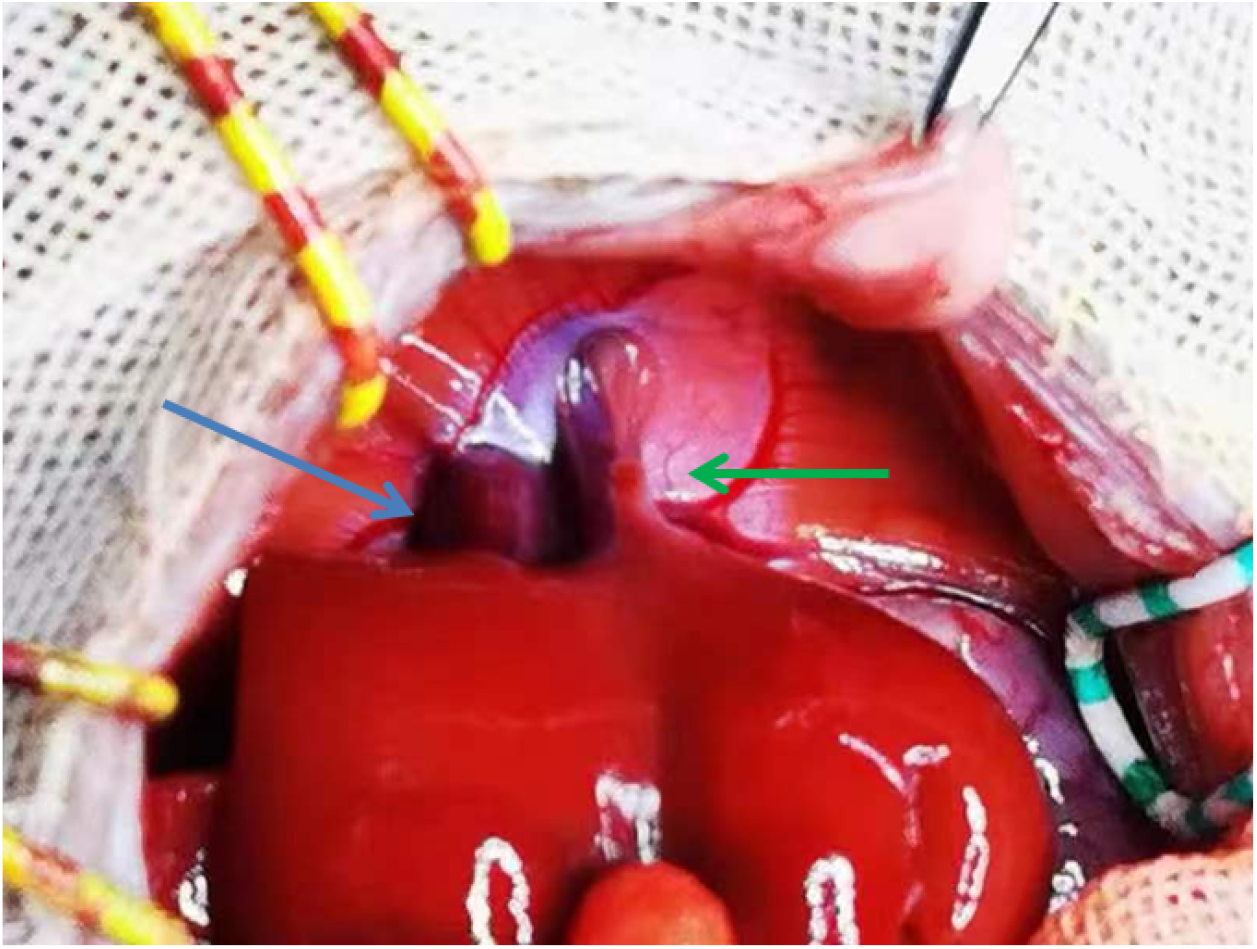
Two outflows in the recipient rat liver. (arrows, green and blue).

**Figure 4.**
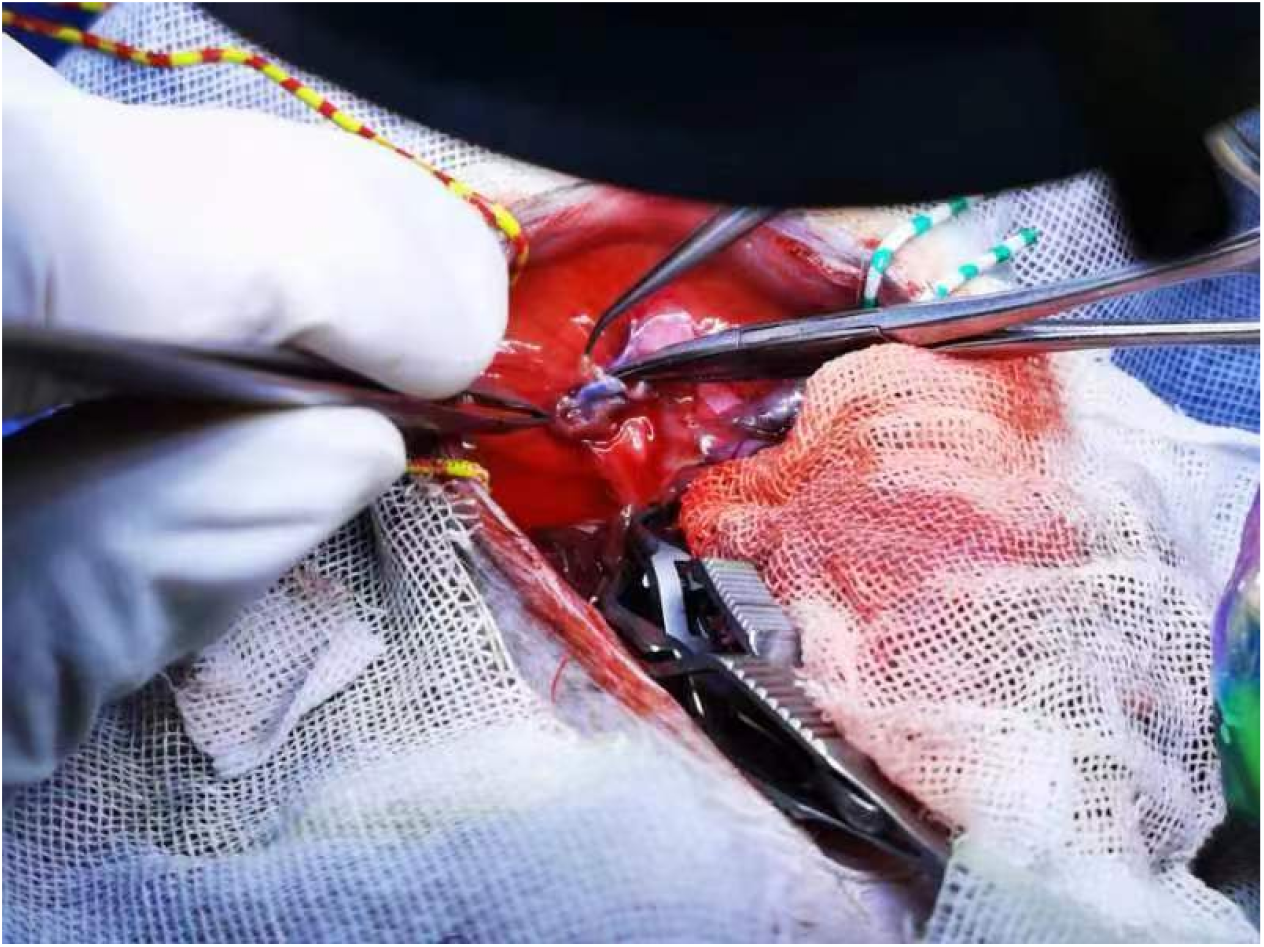
Two outflows of the liver in the recipient rat. The small orifice was closed.

## Discussion

OLT has become well-established and effective treatment exclusively for end-stage liver diseases, benefiting many patients to gain new lives. LT is a complicate surgery and are further needed for basic studies. Small animal liver transplantation models like rats have important value, and the rat still holds a pillar position in the research field of OLT, especially in chronic rejection study [3-6].

Surgically LT is a challenging procedure with dissection and removal of a diseased liver and with subsequent implantation of the graft, effective reconstruction of vascular and biliary anastomoses including potent outflow is fundamentally prerequisite, this requires good condition and normal anatomy of the host including cardiac, pulmonary and renal functionalities [1,2]. Accessory liver lobe (ALL) is a congenital ectopic liver tissue, which is mainly due to embryonic dysplasia described in 1767 [7,8]. Two types of ALL were found: an accessory lobe joining normal liver tissue and the other that is completely separated and often in the thoracic cavity, it remains challenging upon removal of the native liver and has posed problem to OLT. It is un-contemplated clinically and difficult to diagnose non-surgically [9].

Effective hepatic venous drainage is particularly significant in OLT, wider anastomosis is absolutely instrumental. To our knowledge, it is the first report on abnormal outflow of the liver which differs from the Budd-Chiari syndrome that obstruction to hepatic venous drainage occurs, causing ascites, edema, and hepatomegaly [10]; Budd-Chiari syndrome may be due to obstruction without thrombosis, whereas for abnormal outflow, obstruction does not occur before OLT and can occur due to incorrect reconstruction of liver outflow upon OLT. The reason of two outflow orifices of rat liver is unknown. Screening these abnormalities in small animals is also significant in reality.

### Conclusions

Abnormalities or deformed anatomy of the host liver have a negative impact on OLT, suggesting that pre-transplant determination is so significant clinically.

## Abbreviations

OLT,: orthotopic liver transplantation
AHT,: anhepatic time
BN,: Brown Norway
PV,: portal vein
SHVC,: supra-hepatic vena cava
IVC,: infra-hepatic vena cava

## Declarations

### Ethics approval and consent to participate

All experiments (No:HNTCMDW-20240829) were approved by the Ethics Committee of Henan Integrated Traditional Chinese and Western Medicine Hospital, and conducted in compliance with the standards for animal use and care set by ARRIVE guidelines and the Institutional Animal Care Committee of Henan Provincial People’s Hospital and Henan Integrated Traditional Chinese and Western Medicine Hospital, we conducted animal experiments at Henan Integrated Traditional Chinese and Western Medicine Hospital (rental).

### Consent for publication

**N/A**

### Availability of data and materials

All data and materials are available on request,ST Zhou is responsible for all data.

### Competing interests

The authors have no conflicts of interest to declare

### Funding

Purchasing animals, caring and the project of tolerance induction were sponsored by Project 23456 of Henan Provincial People’s Hospital

### Authors’ contributions

YF Chen performed some OLT and revised the draft, Z.Li discussed and collected data. GY. Chen funded the study. ST Zhou performed some OLT, conceived, designed, finalized the study and wrote the draft. All authors have read and approved the manuscript.

## Acknowledgements

**N/A**

## Supplementary Video

**Accessory liver lobe**. Separation was made for small ALL

## Notes

### Competing Interest Statement

The authors have declared no competing interest.

